# Climatic influence on the anthrax niche in warming northern latitudes

**DOI:** 10.1101/219857

**Authors:** Michael G. Walsh, Allard W. de Smalen, Siobhan M. Mor

## Abstract

Climate change is impacting ecosystem structure and function, with potentially drastic downstream effects on human and animal health. Emerging zoonotic diseases are expected to be particularly vulnerable to climate and biodiversity disturbance. Anthrax is an archetypal zoonosis that manifests its most significant burden on vulnerable pastoralist communities. The current study sought to investigate the influence of temperature increases on the landscape suitability of anthrax in the temperate, boreal, and arctic North, where observed climate impact has been rapid. This study also explored the influence of climate relative to more traditional factors, such as livestock distribution, ungulate biodiversity, and soil-water balance, in demarcating high risk landscapes. Machine learning was used to model landscape suitability as the ecological niche of anthrax in northern latitudes. The model identified climate, livestock density and wild ungulate species richness as the most influential landscape features in predicting suitability. These findings highlight the significance of warming temperatures for anthrax ecology in northern latitudes, and suggest potential mitigating effects of interventions targeting megafauna biodiversity conservation in grassland ecosystems, and animal health promotion among small to midsize livestock herds.

**Significance Statement:** We present evidence that a warming climate may be associated with the current distribution of anthrax risk in the temperate, boreal, and arctic North. Moreover, projected warming over the coming decades was associated with substantive expansion of this risk. In addition, livestock distribution, ungulate biodiversity, and soil-water balance were also influential to anthrax risk. While these results are sobering for the future health of livestock and pastoralist communities in the northern latitudes, the coincident modulating effect of ungulate biodiversity may suggest targeted ecosystem conservation as a possible buffer against a growing anthrax niche.

## Introduction

Climate change has begun to manifest significant impact on terrestrial ecosystem structure, particularly in northern latitudes(1). Soil composition, vegetation cover, the distribution of wild ruminant species, and the density and movement of domestic livestock are all shifting as temperatures rise globally and extreme weather events increase(2). As ecosystems are challenged by climatic perturbation, new vulnerabilities may alter infection transmission cycles between hosts and pathogens(3-6). Grasslands in the semi-arid steppe of the northern latitudes are fragile systems that may be particularly vulnerable to the influence of warming (1, 3, 7, 8). This may have important consequences for emerging diseases such as anthrax, as grasslands are important grazing zones for wild ruminants and the domestic livestock of pastoralist communities(9). Indeed, the modifying effects of climate may extend even farther north into the tundra biome, where recent outbreaks of anthrax in the Russian arctic have followed melting permafrost, devastating indigenous caribou-herding communities(10-12).

Anthrax is an important global disease of livestock that carries a high animal mortality and is associated with spillover to humans. Epizootic and zoonotic infection can be particularly impactful to pastoralist communities, where outbreaks exact significant public health, veterinary, and economic consequence(13). Several regions are meso- or hyperenzootic for anthrax in North America, eastern Europe, and central Asia. However, zoonotic transmission is typically well controlled in more affluent countries and less so in resource-poor countries, such that wide global disparity exists in terms of human infection(13).

Anthrax is caused by the spore-forming bacterium, *Bacillus anthracis,* which can remain viable across varied environmental conditions and over long periods of time(13-15). The infection cycle of *B. anthracis* is complex, with the bacterium remaining hidden within the landscape for long periods until suitable environmental conditions are realized. The pathogen prefers alkaline soils, but also occupies a spectrum of pH across the pedosphere (14, 15).The endospores are heat-stable and resistant to desiccation, but exhibit rapid growth with increased precipitation particularly following periods of drought(15, 16). Moreover, the spores are subject to mechanical environmental displacement due to abiotic mechanisms such as flooding events(17-19) and local slope(20), or biotic mechanisms such as tabanid flies(21). With the exception of slope, all such factors could be affected by global warming.

*B. anthracis* may also share a unique relationship with grasses and ruminants. *B.anthracis*-contaminated soil has been shown to enhance the establishment of grass species(22), and these same species demonstrated significantly increased rates of growth in blood-contaminated soils(22, 23). The latter phenomenon is a common occurrence when ruminants succumb to the disease and exude profuse amounts of blood and body fluids into soil following death(13). Thus, anthrax infection ecology may include important interaction between the pathogen, the host, and the host’s herbivory.

The current study sought to investigate the influence of temperature anomalies on the global landscape suitability of anthrax in northern latitudes. A machine learning algorithm was used to model the ecological niche of anthrax outbreaks as a function of climate, soil composition, pasture land use, livestock density and wild ungulate species richness. It was hypothesized that areas of increasing average annual temperature would manifest higher suitability than areas without anomalous warming.

## Methods

Anthrax occurrences were acquired from the World Animal Health Information System (WAHIS) web portal[WAHID], which is the biosurveillance archive maintained by the World Organization for Animal Health (OIE)(24). Each documented occurrence reported the location, date, type of livestock affected, and the number of infected animals. Because the objective of this study was to explore the influence of temperature anomalies on anthrax landscape suitability in the temperate, boreal, and arctic regions of the global North, only outbreaks occurring at or north of 25°N latitude were included. Seventy-six anthrax submissions, reporting on 82 geographically unique outbreaks, were made to OIE between January, 2005 and December, 2016 in this northern extent. Due to a lack of reporting to OIE in the United States (US), Canada, and China, the OIE data were supplemented with additional data obtained from the ProMED-mail electronic surveillance system. This system is maintained by the International Society of Infectious Diseases and provides archival documentation of formal and informal reports of infectious diseases(25). This latter source has been shown to provide good coverage of zoonotic disease events in North America and demonstrates a low rate of reporting error(26). An additional 63 outbreaks were identified in ProMED-mail, two-thirds of which were from the US (n = 17) and Russia (n = 25). The geographic coordinates for all events reported in both OIE and ProMED-mail were obtained in Google Maps as the centroid of the reported municipality. A total of 145 anthrax outbreaks between 2005 and 2016 were available for analysis.

Temperature anomaly data were obtained from the NASA Earth Observation (NEO) data repository for satellite imagery as a 0.5 degree raster for each year from 2005 to 2016(27, 28). Temperature anomaly for any given year between 2005 and 2016 represents the extent of annual temperature divergence (in degrees Celsius) from the homogeneity-adjusted, weighted-average annual temperature during the period 1951-1980(28). The latter period was selected as the baseline, consistent with the detailed study of global temperature change initiated by the Goddard Institute for Space Science (GISS) in 1978. GISS used a three decade period as the baseline climate interval because 30 years are the accepted length of observation required to record one temporal unit of climate(28). This specific period (1951-1980) was used because reliable global temperature estimates were not available prior to 1950(28). In the current investigation, temperature divergence was captured as a global raster for each year under anthrax surveillance (i.e. 2005 to 2016). Mean temperature anomaly was then calculated across the 2005-2016 time period to represent the average warming trend between two serial climatic time points (point 1 = 1951-1980; point 2 = 2005-2016).

In addition to exploring the influence of warming anomalies between the present and the recent past, we also wanted to predict future landscape suitability based on projected future warming. Mean annual temperature projected for the year 2050 was constructed by the Coupled Model Intercomparison Project Phase 5 (CIMP5) and acquired from the WorldClim data repository(29). CIMP5 is a collaborative climate modeling project incorporating twenty modeling groups to predict future climate under multiple representative concentration pathways (RCPs)(30, 31). Projected warming was evaluated at RCPs representing 4.5 W m^-2^ and 8.5 W m^-2^ radiative forcing due to CO^2^ concentrations because these intermediate and high emission scenarios, respectively, have become the standard for modeled climate change projection (32). A raster of projected warming anomalies was used with the estimated Maxent model described above to predict anthrax landscape suitability in 2050.

The Priestley-Taylor *α* coefficient (P-Tα) was used as a robust indicator of water-soil balance in the landscape(33, 34). This coefficient is the ratio of actual evapotranspiration to potential evapotranspiration. It represents water stress in each 1 km^2^ by capturing both water availability in the soil and water requirements of the local vegetation and contrasting this with solar energy input. Thus the measure is a robust estimate of environmental water stress through soil-water balance. A global raster for P-Tα was retrieved from the Consultative Group for International Agricultural Research (CGIAR) Consortium for Spatial Information. The ratio is dimensionless and ranges from 0 (high water stress) to 1 (low water stress)(35).

The Global Soil Dataset for Earth System Modeling was sourced for soil pH and organic content, which is based on an improved protocol of the Harmonized World soil Database(36). The resolution of these two rasters is 5 arc-minutes (approximately 10 km).

Rasters quantifying the proportion of pasture land(37, 38) and mammalian species richness(39) were obtained from the Socioeconomic Data and Applications Center (SEDAC) repository at 5 arc-minutes and 30 arc-seconds, respectively. The latter was used to construct a profile of wild ungluate species richness comprising the Cervidae, Bovidae, Equidae, Camelidae, Antilocapridae, Suidae, and Tayassuidae families. Livestock densities for cattle, sheep, and goats were acquired from the Gridded Livestock of the World (GLW) as 30 arc-second rasters(40).

The human footprint (HFP) was used to weight the sampling of background points to correct for potential reporting bias in anthrax presence points (see modeling description below).The HFP was quantified using data obtained from SEDAC(41). The HFP was calculated in two stages, based on the methods by Sanderson et al. (2002)(42). First, the human influence index (HII) was constructed. The HII measures the impact of human presence on the landscape as a function of eight domains: 1) population density, 2) proximity to railroads, 3) proximity to roads, 4) proximity to navigable rivers, 5) proximity to coastlines, 6) intensity of nighttime artificial light, 7) location in or outside delineated urban space, and 8) land cover. The domains are scored according to the level of human impact per geographic unit, whereby higher scores signify greater human influence. A composite index is then created by combining the eight individual domains. This composite ranges from 0, indicating an absence of human influence (i.e. a parcel of land unaltered by human activity), to 64, indicating maximal human influence in the landscape. The HII composite is subsequently normalized according to the 15 terrestrial biomes defined by the World Wildlife Fund to obtain the HFP. The normalization is represented as a ratio of the range of minimum and maximum HII in each biome to the range of minimum and maximum HII across all biomes, and is expressed as a percentage with a spatial resolution of approximately 1 km^2^ (42).

## Statistical Analysis

Maximum entropy (Maxent) machine learning was used to model the landscape suitability of anthrax across the northern latitudes of the eastern and western hemispheres. Machine learning applications have gained wide application for modeling the ecological niches of many zoonotic infectious diseases(43-45). Maxent in particular is analytically appealing because the model requires no specific functional form. Rather, homogeneity between the outcome and predictors is optimized based on data partition algorithms. Moreover, the locations of unknown (and unknowable) anthrax outbreak absences are not required by the Maxent algorithm to model the system(46, 47). The application of Maxent to ecological niche modeling is a popular implementation due to its robustness(48).

Each of the climate, soil, land cover, livestock density, and wild ungulate species richness features described above were included in the Maxent model. The spatial scale of the analysis was 0.5° of arc, which is approximately 55 km. Therefore the resolution of predicted landscape suitability is 55 km X 55 km. We sampled 10,000 background points, which were weighted by the percentage of the human footprint in the landscape to adjust for potential reporting bias in anthrax surveillance (49). The regularization parameter was selected at 1.0 to correct for overfitting. The Maxent model was trained using five-fold cross-validation, and performance was assessed as the area under the receiver operating characteristic curve (AUC). The AUC was corrected for spatial sorting bias(50). As a sensitivity analysis the anthrax suitability model was repeated at 1° resolution to determine whether landscape suitability was robust to scale. The landscape features were ranked according to their permutation importance in the model, which randomly permutes the values of the factors between background and presence points in the training dataset(46, 52).

Finally, the model was used to predict future landscape suitability based on projected temperature increases using the averaged 2050 CHIMP5 climate models as described above. Future landscape suitability was based on climate projections at the 4.5 and 8.5 RCP scenarios, however these were very similar so we report only the intermediate scenario (4.5 RCP) in the main text and include the high scenario (8.5 RCP) in the supplementary information (S3 Figure 3).

**Figure 3.**
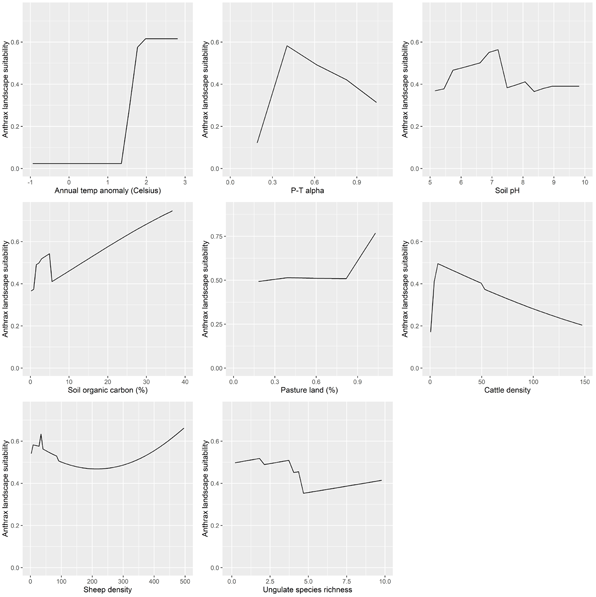
Response curves showing the functional relationships between each feature and anthrax landscape suitability.

The maxent function (dismo package; v. 0.9-3) with Bernoulli distribution was used to fit the model(47, 53, 54). Since this study focused on the northern temperate, boreal, and arctic latitudes, maps of the predicted anthrax niche and each landscape feature were projected to the Lambert azimuthal equal-area to avoid the severe areal distortion in the high northern latitudes associated with other more common projections. All analyses were performed using R statistical software version 3.1.3 (55).

## Results

The distribution of anthrax outbreaks in the temperate, boreal, and arctic regions of the northern hemisphere is presented in Figure 1. The the distribution of the climate, soil and pasture cover, and livestock and wild ungulate species richness features are presented in SI Figure 1.

**Figure 1.**
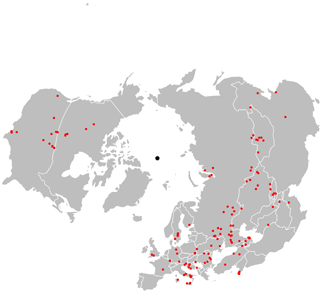
Distribution of anthrax outbreaks as documented by the World Organization for Animal Health (OIE) and Pro-MED mail surveillance mechanisms between 2005 and 2016. For orientation with the Lambert azimuthal equal-area projection of the map, the North Pole is represented as the central black dot.

Climate was highly influential to the landscape suitability of anthrax outbreaks, with warming temperature anomaly (permutation importance = 22.6%) and the P-Tα for water-soil balance (permutation importance = 25.8%) together contributing 48.4% in permutation importance (Figure 2). Cattle density (21.6%) and wild ungulate species richness (9.1%) were also important delineators of landscape suitability. While not dominating features soil pH, organic carbon content, and sheep density nevertheless were still impactful in the model, collectively contributing 16.8% to the loss function. Model performance was reasonable with the AUC, adjusted for spatial sorting bias, equal to 78%.

**Figure 2.**
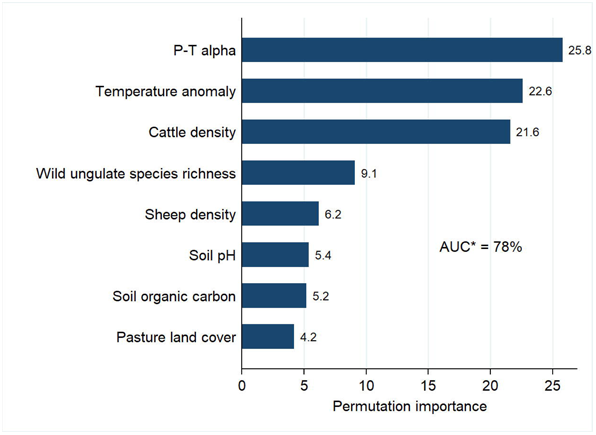
Landscape feature ranking according to each feature's permutation importance in the Maxent model. The area under the curve (AUC) is reported as a percentage and adjusted (*) for spatial sorting bias.

The response curves showing the functional effect of each landscape feature conditional on all others are presented in Figure 3. Landscape suitability increased dramatically in areas of large anomalous warming. Suitability was high in the areas of mild to moderate water stress with P-Tα peaking at 0.37, which is typical of semi-arid steppe landscapes. However, landscape suitability was low in areas of extreme water stress or in areas absent of water stress. Soil pH in the range of 5 - 7.5, and soils with higher organic carbon content, were associated with greater landscape suitability, as was the presence of pasture, particularly in high proportion. Cattle and sheep density were both influential in the model, however cattle density was associated with greatest risk among smaller herds while sheep demonstrated elevated risk at low and high density. Finally, increasing wild ungulate species richness was associated decreasing anthrax suitability.

The predicted landscape suitability of anthrax outbreaks in the northern latitudes is presented in Figure 4 (left panel). In the eastern hemisphere, an intercontinental anthrax suitability corridor emerged that extended from Eastern and Mediterranean Europe across the steppe of Russia and central Asia and into northwestern China. In addition, two narrow high risk corridors extended from the intercontinental anthrax belt southward down the eastern and western boundaries of central Asia. There were also prominent areas of high landscape suitability in some areas of the Arctic and sub-Arctic North, particularly in northern Siberia, Iceland, and parts of Scandinavia and the UK, although these were not as extensive as the steppe regions to the south. A marked corridor of landscape suitability also emerged in the western hemisphere, running from north to south, and corresponding to the landscape configuration of steppe country in North America. The areas of greatest risk were in south-central Canada and the north-central United States. However, there were some areas of high landscape suitability identified in the high Arctic. The sensitivity analysis showed no meaningful differences in landscape suitability when evaluated at 1° resolution (S2 Figure 2). Landscape suitability predicted by 2050 identified the same general regions of the temperate, boreal, and arctic North, but risk had increased and expanded substantively in many areas, particularly in North America, southern Central Asia, Mediterranean Europe, and China (Figure 4; right panel).

**Figure 4.**
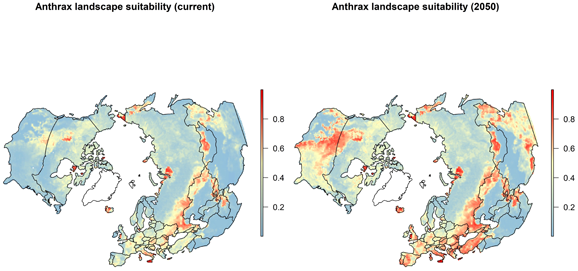
Predicted anthrax landscape suitability based on current warming anomalies (left panel), and future landscape suitability based on projected anomalies in 2050 using a representative concentration pathway of 4.5 W m^-2^ radiative forcing due to CO^2^ concentration (right panel). Greenland is shown in outline only as there was insufficient data for each landscape feature to model the suitability.

## Discussion

This investigation is the first to describe the landscape suitability of anthrax across the temperate, boreal, and arctic North of both the eastern and western hemispheres. We found that climate and the distribution of domestic livestock and wild ungulate species richness were important delineators of suitability. Warming annual temperature anomalies were associated with increasing suitability, as was a water-soil balance that favored mild to moderate water stress. This is the first study to show an increase in anthrax risk across the northern latitudes associated with a warming climate, an association that was highly influential to the predicted niche. Moreover, landscape suitability is predicted to expand and increase within a relatively short period of time (~30 years) with continued warming. The effects of livestock densities and soil composition notwithstanding, these results highlight the potential effects of rising temperatures and moderate water stress in the landscape on anthrax risk in northern latitudes. In addition, increasing wild ungulate species richness was associated with diminishing suitability and may indicate that megafauna biodiversity protects against anthrax transmission, or this biodiversity may simply be a proxy for less pathogenic ecosystems. While this is the first study to explore anthrax suitability across extensive northern latitudes, our results identified a niche that was generally in geographical agreement with other regional ecological niche studies(56-58).The ecological niche identified by the current study corroborated the findings of two previous studies in the United States(56) and Kazakhstan(57), although landscape suitability predicted for Texas in the current study was not as strong as that predicted in the previous United States niche model. Both of these earlier studies also included measures of climate in their models, however only the latter incorporated measures of climate change. A third study of anthrax risk in China was less convergent with our current findings(59). The two studies did agree on predicted risk with respect to the northeastern and northwestern regions of China, however they diverged on the predicted risk for central China. These differences are not surprising given that the previous study used surveillance data aggregated at the county and province level and also generalized several disparate measures of climate.

In the current study, areas where recent annual temperatures (2005 to 2016) exhibited warming divergence from past annual temperatures (1950 to 1980) were more suitable to anthrax outbreaks than those with no change or cooling divergence. These areas of anomalous warming were most pronounced in the higher latitudes. There are several mechanisms by which warming may influence suitability across intercontinental scales of varying ecosystem structure. First, warming trends in the Arctic have been associated with substantial permafrost melt(60-62), which may operate directly in the infection cycle by thawing wild or domestic ruminant carcasses, or their previously frozen tissues or body fluids, and subsequently releasing spores into the soil and activating new growth(11, 63). This phenomenon has been documented at the animal burial sites of previous epizootics(10, 12, 64). Second, warming in semi-arid steppe or shrubland may increase dry conditions that can be favorable to anthrax spore preservation, dissemination and activation in the middle and southerly northern latitudes, particularly when summer droughts are punctuated with intermittent precipitation(17-19). The current study lends further support to this as landscape suitability appeared to be optimized in areas with a mild to moderate degree of water stress, which is typical of the grasslands and steppe of eastern Europe, north-central Asia and north-central North America(65). Furthermore, dry soils may favor mechanical dissemination of spores by intermittent precipitation events (66). Third, hot weather may modify host susceptibility by decreasing immunogenicity(67, 68). Therefore, warming temperatures and dry conditions may enhance the transmission cycle of *B. anthracis* by simultaneously opening new chains of transmission, promoting mechanical motility through the environment, or suppressing host immune function. While the climatic features in the current study support an infection ecology modulated by warming annual temperatures and moderate soil-water stress, none of the potential mechanisms described above were directly observed in the current study and, as such, definitive pathways describing anthrax epidemiology must await interrogation by field studies of livestock, wildlife, and the environment.

The distribution of animals in the landscape was influential to anthrax suitability, with livestock density and wild ungulate species richness accounting for one third (36.9%) combined permutation importance. Small herds were associated with greatest risk among cattle, while sheep delineated anthrax suitability in low and high density. This may reflect greater adoption of preventive measures (i.e. vaccination) in areas supporting large, or industrial, cattle industries(69, 70), whereas such prevention may be more lacking in general in sheep husbandry. Increasing ungulate species richness decreased landscape suitability for anthrax, which is consistent with the view that high biodiversity has a diluting effect on pathogen transmission (71, 72)

Soil organic carbon and pH have long been recognized as important properties in the infection ecology and transmission cycle of *B. anthracis*(14, 15). High organic carbon, and pH in the range of 6-7, delineate the Mollisol and Chernozem soils of the prairies of North America and the steppe of eastern Europe and central Asia, respectively, which are typically favorable to anthrax sporulation in northern latitudes(15, 66). These soils tend to be rich in calcium, which can further promote spore germination under the right environmental conditions(14, 73, 74). Moreover, these soil profiles support rich grassland ecosystems and agricultural production(75), which is likely reflected in the influence of both pasture land cover and livestock densities on anthrax landscape suitability. Thus, while soil composition was not as influential as climate and livestock density, the current study does reinforce the pedological and edaphological paradigm for *B. anthracis* across both the western and eastern hemispheres.

This study has some important limitations. First, the documented anthrax outbreaks included in this study are derived from the OIE and ProMED surveillance systems, which may not have captured all anthrax outbreak occurrences and which may be more responsive in areas with better reporting infrastructure, thus potentially leading to reporting bias in the identified anthrax locations. We attempted to correct for such reporting bias by selecting background points weighted by the presence of the human footprint as a proxy for reporting infrastructure. Nevertheless, we must concede that these data may not be representative of the total anthrax outbreaks occurring in northern latitudes over the time period under study. Second, the scale of the study is coarse. While this is unlikely to be of substantial influence to landscape features operating at small (i.e. coarse) spatial scale, such as temperature, soil composition, and wildlife species range, it would most certainly be influential to features operating at large (i.e. fine) scale, such as subtle changes in topographic slope and water drainage, or the distribution of tabanid flies as mechanical vectors. As such, we were not able to investigate the additional influences of hyper-local features such as water pooling, which has been previously identified as an important delimiter of anthrax risk due to spore accumulation through water dispersal(20), or the ecological niche of tabanids despite convincing evidence that these may be important to anthrax transmission in some regions(21). Third, temperature anomaly was assessed over a relatively recent time frame (i.e. change between 1950-1980 and 2005-2016), and therefore assumes as its baseline a temperature regime that was already likely undergoing warming trends. Nevertheless, we would expect this to dilute the effect of global warming on anthrax landscape suitability. However, given the strong influence that was still apparent in the current analysis, it would seem that temperature increase is impactful to the anthrax niche even in the short term.

In conclusion, this investigation revealed the novel finding that, mean global temperature increases between 1950-1980 and 2005-2016 identified high landscape suitability for anthrax outbreaks in the northern latitudes during the early decades of the twenty-first century. Continued warming in northern latitudes predicted an expanding anthrax niche. In addition, livestock density and wild ungulate species richness were also strongly associated with anthrax suitability. These results provide the first evidence that climate change may influence the risk of this impactful zoonotic disease across a large extent of the northern latitudes. Additional findings from this study suggest that interventions targeting megafauna biodiversity conservation in grassland ecosystems, and animal health promotion among small to midsize livestock herds (e.g. vaccine promotion among pastoralist communities) may reduce the overall landscape suitability to future outbreaks, possibly mitigating some of the effects of climate change.

## References

1. Walther G-R (2010) Community and ecosystem responses to recent climate change. Philos Trans RSoc London B Biol Sci 365(1549). Available at: http://rstb.royalsocietypublishing.org/content/365/1549/2019.short [Accessed July 11, 2017].

2. IPCC (2014) Impacts, adaptation, and vulnerability. Part A: global and sectoral aspects. Contribution of Working Group II to the fi fth assessment report of the Intergovernmental Panel on Climate Change. Climate Change, eds Field C, Barros V, Dokken D (Cambridge University Press, Cambridge).

3. Hoberg EP, Polley L, Jenkins EJ, Kutz SJ (2008) Pathogens of domestic and free-ranging ungulates: global climate change in temperate to boreal latitudes across North America. Rev Sci Tech 27(2):511–28.

4. Hoberg E, Brooks DR (2015) Evolution in action: climate change, biodiversity dynamics and emerging infectious disease. Philos Trans R Soc B Biol Sci 370(1665):20130553–20130553.

5. Parkinson AJ, et al. (2014) Climate change and infectious diseases in the Arctic: establishment of a circumpolar working group. Int J Circumpolar Health 73(1):25163.

6. Whitmee S, et al. (2015) Safeguarding human health in the Anthropocene epoch: report of The Rockefeller Foundation–Lancet Commission on planetary health. Lancet 386(10007):1973–2028.

7. Hopkins A, Del Prado A (2007) Implications of climate change for grassland in Europe: impacts, adaptations and mitigation options: a review. Grass Forage Sci 62(2):118–126.

8. Soussana J-F, Lüscher A (2007) Temperate grasslands and global atmospheric change: a review. Grass Forage Sci 62(2):127–134.

9. Neely C dq (Constance), Bunning S, Wilkes A (2009) Review of evidence on drylands pastoral systems and climate changed: implications and opportunities for mitigation and adaptation (Food and Agriculture Organization of the United nations (FAO)) Available at: https://books.google.com.au/books?id=9TFAtwAACAAJ&dq=Review+of+evidence+on+drylands+pastoral+systems+and+climate+change&hben&sa=X&redir_esc=y [Accessed July 11, 2017],

10. Revich B, Podolnaya MA (2011) Thawing of permafrost may disturb historic cattle burial grounds in East Siberia. Glob Health Action 4(1):8482.

11. Revich B, Tokarevich N, Parkinson AJ (2012) Climate change and zoonotic infections in the Russian Arctic. Int J Circumpolar Health 71:18792.

12. Elvander M, Persson B, Sternberg Lewerin S (2017) Historical cases of anthrax in Sweden 1916-1961. Transbound Emerg Dis 64(3):892–898.

13. International Office of Epizootics., World Health Organization., Food and Agriculture Organization of the United Nations. (2008) Anthrax in humans and animals. (World Health Organization).

14. Dragon D, Rennie RP (1995) The ecology of anthrax spores: tough but not invincible. Can VetJ-La Rev Vet Can 36(5):295–301.

15. Hugh-Jones M, Blackburn J (2009) The ecology of Bacillus anthracis. Mol Aspects Med 30(6):356–367.

16. Anthrax in Humans and Animals (2008) (World Health Organization) Available at: http://www.ncbi.nlm.nih.gov/pubmed/26269867 [Accessed July 10, 2017].

17. Dragon D, Elkin B, Nishi J, Ellsworth TR (1999) A review of anthrax in Canada and implications for research on the disease in northern bison. J Appl Microbiol 87(2): 208–13.

18. Fox MD, Boyce JM, Kaufmann AF, Young JB, Whitford HW (1977) An epizootiologic study of anthrax in Falls County, Texas. J Am Vet Med Assoc 170(3):327–33.

19. Gaitanis G, et al. (2016) An Aggregate of Four Anthrax Cases during the Dry Summer of 2011 in Epirus, Greece. Dermatology 232(1):112–6.

20. Van Ness GB (1971) Ecology of anthrax. Science 172(3990):1303–7.

21. Blackburn J, Van Ert M, Mullins J, Hadfield T, Hugh-Jones ME (2014) The Necrophagous Fly Anthrax Transmission Pathway: Empirical and Genetic Evidence from Wildlife Epizootics. Vector-Borne Zoonotic Dis 14(8):576–583.

22. Turner WC, et al. (2014) Fatal attraction: vegetation responses to nutrient inputs attract herbivores to infectious anthrax carcass sites. ProcRSoc B Biol Sci 281(1795):20141785–20141785.

23. Ganz HH, et al. (2014) Interactions between Bacillus anthracis and Plants May Promote Anthrax Transmission. PLoS Negl Trop Dis 8(6):e2903.

24. Health WO for A World Animal Health Information System. Available at: http://www.oie.int/wahis_2/public/wahid.php/Diseaseinformation/lmmsummary.

25. International Society for Infectious Diseases ProMED-mail. Available at: http://www.promedmail.org/.

26. Cowen P, et al. (2006) Evaluation of ProMED-mail as an electronic early warning system for emerging animal diseases: 1996 to 2004. J Am Vet Med Assoc 229(7): 1090–1099.

27. NASA Earth Observations Available at: https://neo.sci.gsfc.nasa.gov/view.php?datasetld=GISS_TA_M.

28. Hansen J, Ruedy R, Sato M, Lo K (2010) GLOBAL SURFACE TEMPERATURE CHANGE. Rev Geophys 48(4):RG4004.

29. WorldClim WorldClim - Future Climate Data. Available at: http://www.worldclim.org/CMIP5vl.

30. Programme WCR (2011) WCRP Coupled Model Intercomparison Project - Phase 5. CLIVAR Exch 15(56).

31. Taylor KE, et al. (2012) An Overview of CMIP5 and the Experiment Design. Bull Am Meteorol Soc 93(4):485–498.

32. Moss RH, et al. (2010) The next generation of scenarios for climate change research and assessment. Nature 463(7282):747–56.

33. Priestley CHB, Taylor RJ (1972) On the Assessment of Surface Heat Flux and Evaporation Using Large-Scale Parameters. Mon Weather Rev 100(2):81–92.

34. Khaldi a., Khaldi a., Hamimed a. (2014) Using the Priestley-Taylor expression for estimating? actual évapotranspiration from satellite Landsat ETM + data. Proc Int Assoc Hydrol Sci 364(June):398–403.

35. Trabucco A, Zomer RJ (2010) Global Soil Water Balance Geospatial Database. CGIAR Consort Spat Inf. Available at: http://www.cgiar-csi.org.

36. Shangguan W, Dai Y, Duan Q, Liu B, Yuan H (2014) A global soil data set for earth system modeling. JAdv Model Earth Syst 6(1):249–263.

37. Ramankutty N, Evan AT, Monfreda C, Foley JA (2010) Global Agricultural Lands: Pastures, 2000. doi:10.7927/H47H1GGR.

38. Ramankutty N, Evan AT, Monfreda C, Foley JA (2008) Farming the planet: 1. Geographic distribution of global agricultural lands in the year 2000. Global Biogeochem Cycles 22(1):n/a-n/a.

39. Socioeconomic Data and Applications Center | SEDAC Global Mammal Richness Grids. Available at: http://sedac.ciesin.columbia.edu/data/set/species-global-mammal-richness-2015.

40. Robinson TP, et al. (2014) Mapping the global distribution of livestock. PLoS One 9(5):e96084.

41. Socioeconomic Data and Applications Center | SEDAC MethodsEB» Last of the Wild, v2 | SEDAC. Available at: http://sedac.ciesin.columbia.edu/data/collection/wildareas-v2/methods [Accessed December 23, 2014],

42. Sanderson EW, et al. (2002) The Human Footprint and the Last of the Wild. 52(10).

43. Peterson AT (2014) Mapping Disease Transmission Risk: Enriching Models Using Biogeography and Ecology (Johns Hopkins University Press, Baltimore).

44. Hay SI, et al. (2013) Global mapping of infectious disease. Philos Trans R Soc 368(1614):20120250.

45. Stevens KB, Pfeiffer DU (2011) Spatial modelling of disease using data- and knowledge-driven approaches. Spat Spatiotemporal Epidemiol 2(3):125–133.

46. Phillips SJ, Anderson RP, Schapire RE (2006) Maximum entropy modeling of species geographic distributions. Ecol Modell 190(3-4):231–259.

47. Franklin J (2010) Mapping Species Distributions: Spatial Inference and Prediction (Cambridge University Press, Cambridge). First Available at: https://books.google.com/books?id=CkshAwAAQBAJ&pgis=l [Accessed June 8, 2015].

48. Duan R-Y, Kong X-Q, Huang M-Y, Fan W-Y, Wang Z-G (2014) The predictive performance and stability of six species distribution models. PLoS One 9(11):e112764.

49. Phillips SJ, et al. (2009) Sample selection bias and presence-only distribution modelsBI: implications for background and pseudo-absence data Reference Sample selection bias and presence-only distribution modelsBI: implications for background and pseudo-absence data. 19(1):181–197.

50. Hijmans RJ, Hall W (2016) Cross-validation of species distribution modelsBI: removing spatial sorting bias and calibration with a null model Published bySl: Ecological Society of America Stable URLE3: http://www.jstor.org/stable/23143954 REFERENCES Linked references are available on. Ecology 93(3):679–688.

51. Raes N, Ter Steege H (2007) A null-model for significance testing of presence-only species distribution models. Ecogrophy (Cop) 30(5):727–736.

52. Phillips SJ, Dudík M (2008) Modeling of species distribution with Maxent: new extensions and a comprehensive evalutation. Ecograpy 31(December 2007):161–175.

53. Hijmans RJ, Phillips S, Leathwick JR, Elith J (2014) Package "dismo." Compr R Arch Netw. 1–65.

54. Fielding AH, Bell JF (1997) A review of methods for the assessment of prediction errors in conservation presence/absence models. Environ Conserv 24:38–49.

55. Team RC (2016) R: A language and environment for statistical computing. Available at: https://www.r-project.org/.

56. Blackburn JK, McNyset KM, Curtis A, Hugh-Jones ME (2007) Modeling the geographic distribution of Bacillus anthracis, the causative agent of anthrax disease, for the contiguous United States using predictive ecological [corrected] niche modeling. Am J Trop Med Hyg 77(6):1103–10.

57. Joyner TA, et al. (2010) Modeling the Potential Distribution of Bacillus anthracis under Multiple Climate Change Scenarios for Kazakhstan. PLoS One 5(3):e9596.

58. Mullins JC, et al. (2013) Ecological Niche Modeling of Bacillus anthracis on Three Continents: Evidence for Genetic-Ecological Divergence? PLoS One 8(8):e72451.

59. Chen W-J, et al. (2016) Mapping the Distribution of Anthrax in Mainland China, 2005–2013. PLoS Negl Trop Dis 10(4):e0004637.

60. Osterkamp TE, Romanovsky VE (1999) Evidence for warming and thawing of discontinuous permafrost in Alaska. Permafr Periglac Process 10(1):17–37.

61. Stendel M, Christensen JH (2002) Impact of global warming on permafrost conditions in a coupled GCM. Geophys Res Lett 29(13):1632.

62. Davidson EA, Janssens IA (2006) Temperature sensitivity of soil carbon decomposition and feedbacks to climate change. Nature 440(7081):165–173.

63. Dragon DC, Bader DE, Mitchell J, Woollen N (2005) Natural Dissemination of Bacillus anthracis Spores in Northern Canada. Appl Environ Microbiol 71(3):1610–1615.

64. Goudarzi S (2016) What Lies Beneath. Sci Am 315(5):11–12.

65. Sala OE, Lauenroth WK (1982) Small rainfall events: An ecological role in semiarid regions. Oecologia 53(3):301–304.

66. Wall DH, Nielsen UN, Six J (2015) Soil biodiversity and human health. Nature 528(7580):69–76.

67. World Meteorological Organization (1989) Hot Weather Stress. Animal Health and Production at Extremes of WeatherlA: Reports of the CAgM Working Groups on Weather and Animal Disease and Weather and Animal Health. (Secretariat of the World Meteorological Organization), pp 61–122.

68. Webster AJ (1981) Weather and infectious disease in cattle. Vet Rec 108(9):183–7.

69. Bengis RG, Frean J (2014) Anthrax as an example of the One Health concept. Rev Sci Tech 33(2):593–604.

70. Kracalik I, et al. (2014) Changing Patterns of Human Anthrax in Azerbaijan during the Post-Soviet and Preemptive Livestock Vaccination Eras. PLoS Negl Trop Dis 8(7):e2985.

71. Keesing F, et al. (2010) Impacts of biodiversity on the emergence and transmission of infectious diseases. Nature 468(7324):647–652.

72. Civitello DJ, et al. (2015) Biodiversity inhibits parasites: Broad evidence for the dilution effect. Proc Natl Acad Sci 112(28):8667–8671.

73. Foerster HF, Foster JW (1966) Endotrophic calcium, strontium, and barium spores of Bacillus megaterium and Bacillus cereus. j Bacteriol 91(3): 1333–45.

74. Stastná J, Vinter V (1970) Spores of microorganisms. 23. Interdependence of intra- and extracellular levels of calcium: its effect on the germination of bacterial spores in different media. Folia Microbiol (Praha) 15(2):103–10.

75. Eswaran H, Van Den Berg E, Reich P (1993) Organic Carbon in Soils of the World. Soil Sci Soc Am J 57(1):192.

